# Mimicking the breast metastatic microenvironment: characterization of a novel syngeneic model of HER2^+^ breast cancer

**DOI:** 10.1101/2024.01.25.577282

**Authors:** Aaron G. Baugh, Edgar Gonzalez, Valerie H. Narumi, Jesse Kreger, Yingtong Liu, Christine Rafie, Sofi Castanon, Julie Jang, Luciane T. Kagohara, Dimitra P. Anastasiadou, James Leatherman, Todd D. Armstrong, Isaac Chan, George S. Karagiannis, Elizabeth M. Jaffee, Adam MacLean, Evanthia T. Roussos Torres

## Abstract

Preclinical murine models in which primary tumors spontaneously metastasize to distant organs are valuable tools to study metastatic progression and novel cancer treatment combinations. Here, we characterize a novel syngeneic murine breast tumor cell line, NT2.5-lung metastasis (- LM), that provides a model of spontaneously metastatic neu-expressing breast cancer with quicker onset of widespread metastases after orthotopic mammary implantation in immune-competent NeuN mice. Within one week of orthotopic implantation of NT2.5-LM in NeuN mice, distant metastases can be observed in the lungs. Within four weeks, metastases are also observed in the bones, spleen, colon, and liver. Metastases are rapidly growing, proliferative, and responsive to HER2-directed therapy. We demonstrate altered expression of markers of epithelial-to-mesenchymal transition (EMT) and enrichment in EMT-regulating pathways, suggestive of their enhanced metastatic potential. The new NT2.5-LM model provides more rapid and spontaneous development of widespread metastases. Besides investigating mechanisms of metastatic progression, this new model may be used for the rationalized development of novel therapeutic interventions and assessment of therapeutic responses targeting distant visceral metastases

**SUMMARY STATEMENT:** We characterize a new syngeneic, immune-competent murine model of breast cancer (NT2.5-LM) that yields rapid and widespread metastases, preserves spontaneous metastasis, and provides a model for studying novel therapeutic interventions.

## INTRODUCTION

Breast cancer remains one of the leading causes of cancer mortality among women worldwide, with metastatic burden as the major contributor of patient death.(Riggio et al., 2020; Sung et al., 2021) The development of murine models of breast cancer has provided researchers with the means to more intricately study tumor initiation, progression, metastasis, and response to therapies, leading to our current understanding of the complex physiological systems and molecular mechanisms underlying these processes.(Kim and Baek, 2010; Park et al., 2018) Various transgenic models of breast cancer that develop spontaneous mammary tumors and metastases exist.(Green et al., 2000; Chantale T Guy et al., 1992; C T Guy et al., 1992; Lin et al., 2004; Macleod and Jacks, 1999; Siegel et al., 2003) However, only few of these models allow for efficient study of the metastatic tumor microenvironment (TME). Syngeneic models of breast cancer, which involve orthotopic implantation of tumor cells or tumor chunks, are widely utilized, but often times, these models are either slow-growing or do not develop clinically overt metastases. Experimental metastasis models, which involve tail vein injection of tumor cells, are also widely utilized, but these models are limited by lack of resolution in metastatic progression, and conclusions drawn from these models may be artificial. As such, development of appropriate mouse models of breast carcinoma that recapitulate metastatic progression in a pathophysiological and clinically relevant context is necessary.

The immunotolerant MMTV-HER2/Neu (ERBB2) transgenic murine model (NeuN) originally characterized by Guy et al.,(C T Guy et al., 1992) in which FVB/N strain mice express the non-transforming rat *Neu* cDNA under control by a mammary tissue-specific promoter, gives rise to spontaneous mammary tumors between 125 and 300 days. This model yields spontaneously developing mammary tumors that closely mimic human epidermal growth factor 2-positive (HER2^+^) tumors.(Fry et al., 2017) One caveat of this model is its long latency for development of both primary and metastatic disease, as well as the lack of penetrance of metastatic disease. To circumvent these issues, previous efforts have focused on its improvement and have led to the development of a syngeneic tumor cell line derivative, known as NT2.5. The latter model has significantly shortened the time from tumor cell injection to tumor growth and is capable of establishing widespread distant metastases upon cardiac or tail vein injections.(R Todd Reilly et al., 2000; Song et al., 2008) Metastases in various organs can be observed within 3 weeks of NT2.5 tumor cell injection, but this model is also limited by its inability to recapitulate the process of spontaneous metastasis.

In this study, we report the serial passaging of the original NT2.5 cell line to generate a new subline called NT2.5-LM, which represents an orthotopic, immunotolerant model of HER2^+^ breast cancer capable of promoting development of spontaneous metastases. We also perform an in-depth characterization of the newly established NT2.5-LM cell line at both the genomic and proteomic levels to establish the foundations for its potential use in preclinical studies.

## RESULTS

### Orthotopic implantation of NT2.5-LM leads to decreased survival, larger mammary tumors, and increased lung metastasis

In the NT2.5 syngeneic model, NT2.5 cells are implanted in the mammary fat pad of adult female NeuN mice, after which the maximum allowable volume of 1.5 cm^3^ is reached in 4-5 weeks,(Brian J. Christmas et al., 2018; R T Reilly et al., 2000a; Sidiropoulos et al., 2022) prior to the establishment of metastatic disease and preventing efficient study of metastatic tumor microenvironments (TMEs). To derive a highly metastatic cell line, lung metastases were macro-dissected from the lungs of NT2.5 mammary tumor-bearing NeuN mice, dissociated to single-cell suspensions, and intravenously injected into non-tumor-bearing NeuN mice, after which lung metastases were harvested again and the process repeated. After the third round of harvest, spontaneous lung metastases could be observed 3 weeks following mammary fat pad injection of isolated cells, thus establishing the NT2.5-lung metastasis (-LM) cell line for use.

To characterize the phenotype of NT2.5-LM-derived tumors *in vivo*, we orthotopically injected NT2.5-LM cells into the mammary fat pad of NeuN mice and measured survival, tumor burden, and metastatic burden. When compared to parental NT2.5 controls, mice orthotopically injected with NT2.5-LM cells experienced significantly decreased survival (**Fig. 1A**) and increased weekly mammary tumor growth rates (**Fig. 1B**). Despite surgical resection of NT2.5-LM mammary tumors at 12 days post-injection, tumors regrew at 24 days post-injection and reached endpoint criteria faster than NT2.5 mammary tumors (**Figs. S1A-B**). Necropsy analyses of mice with NT2.5-LM mammary tumors revealed widespread metastases in the heart, lymph nodes, lungs, kidneys, adrenal glands, stomach, colon, spleen, skull, ears, body walls, and teeth (**Fig. S2)**, with high metastatic burden observed in the lungs. Moving forward, we focused on the lungs as a surrogate measure of total metastatic burden. When examining lungs of mice euthanized from 34 to 41 days post-injection, we found a significant increase in the number of lung metastases in the NT2.5-LM model, when compared to the NT2.5 control **(Fig. 1C**). NT2.5-LM lung micro-metastases could be observed by H&E staining as early as 7 days post-injection, with consistent growth observed at 10, 22, 28, and 35 days post-injection (**Fig. 1D**).

**Figure 1:**
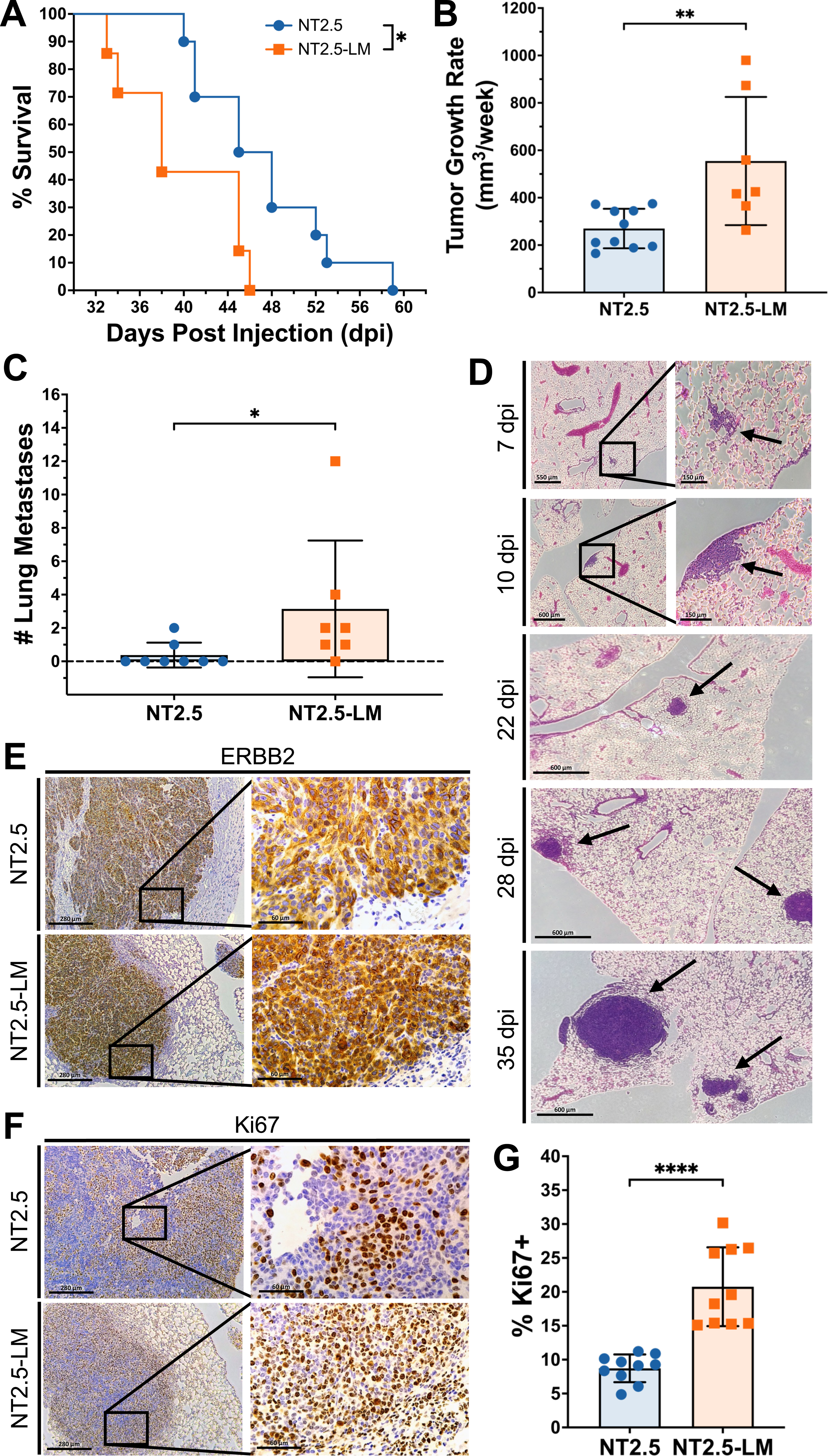
NT2.5-LM leads to decreased survival, larger mammary tumors, and increased lung metastasis. **(A**) 1x10^5^ NT2.5 or NT2.5-LM cells were injected into the mammary fat pad of NeuN mice (NT2.5, n=10; NT2.5-LM, n=7). After surgical resection of NT2.5-LM tumor-bearing mice at 12 days post-injection (dpi), mice were allowed to reach humane survival endpoint with tumor volume exceeding 1.5 cm^3^. (**B)** Mammary tumor sizes of mice in (A) were measured at least 3x a week by calipers, averaged, and used to calculate differences in average weekly tumor growth rate. **(C)** At survival endpoint of mice in (A), the number of surface metastases was counted by visual inspection. **(D)** H&E staining of lungs in NT2.5-LM tumor-bearing mice collected at 7, 10, 22, 28, and 35 days post-injection (dpi). Black arrows point to lung metastases. Scale bars as shown. **(E)** Immunohistochemistry (IHC) staining of Erbb2 and **(F)** Ki67 in NT2.5 mammary tumors (top) and NT2.5-LM lung metastases (bottom) collected at 35 days post-injection. Scale bars as shown. **(G)** Percentage of Ki67+ cells from 10 regions of interest (ROIs) were counted from Ki67 IHC staining in (F). Statistics used: Mantel-Cox Log-rank test for (A), Mann-Whitney U-test for (B-D), Welch’s T-test for (G), *p < 0.05, **p < 0.01, ****p < 0.0001.

To further illuminate on the phenotypic characteristics of NT2.5-LM metastases, we performed immunohistochemical staining for ERBB2, Ki67, CK5, CK6, AE1/3, and EGFR. NT2.5-LM lung metastases are ERBB2-positive (**Fig. 1E**), express similarly low levels of AE1/3 and EGFR, and are similarly negative for CK5 and CK6, when compared to NT2.5 mammary tumors (**Fig. S3**). Finally, NT2.5-LM lung metastases are more proliferative, as observed by increased numbers of Ki67+ cells (**Figs. 1F-G**).

### NT2.5-LM responds to HER2 directed therapy

Patients with HER2^+^ breast cancer demonstrate a response rate of over 35% when treated with HER2-directed monoclonal antibody therapy.(Vogel et al., 2002) To characterize the sensitivity of the NT2.5-LM model to a similar type of therapy, NT2.5-LM metastasis-bearing mice were treated with anti-HER2 antibody by intraperitoneal (i.p.) injection once a week and assessed for survival (**Fig. S4**). Anti-HER2-treated mice showed improved survival when compared to vehicle-treated mice, with a ∼35% response rate to therapy (**Fig. 2A**), similar to that observed in patients treated with single agent therapy.(Vogel et al., 2002) When assessing the anti-HER2 treatment effects on lung metastases, we found that treatment did not change the number of lung metastases (**Fig. 2B**), but it significantly decreased the area of metastases within the lung (**Fig. 2C**). Together, these data suggest that the new NT2.5-LM model demonstrates clinical relevance with regards to its therapeutic response to anti-HER treatments.

**Figure 2:**
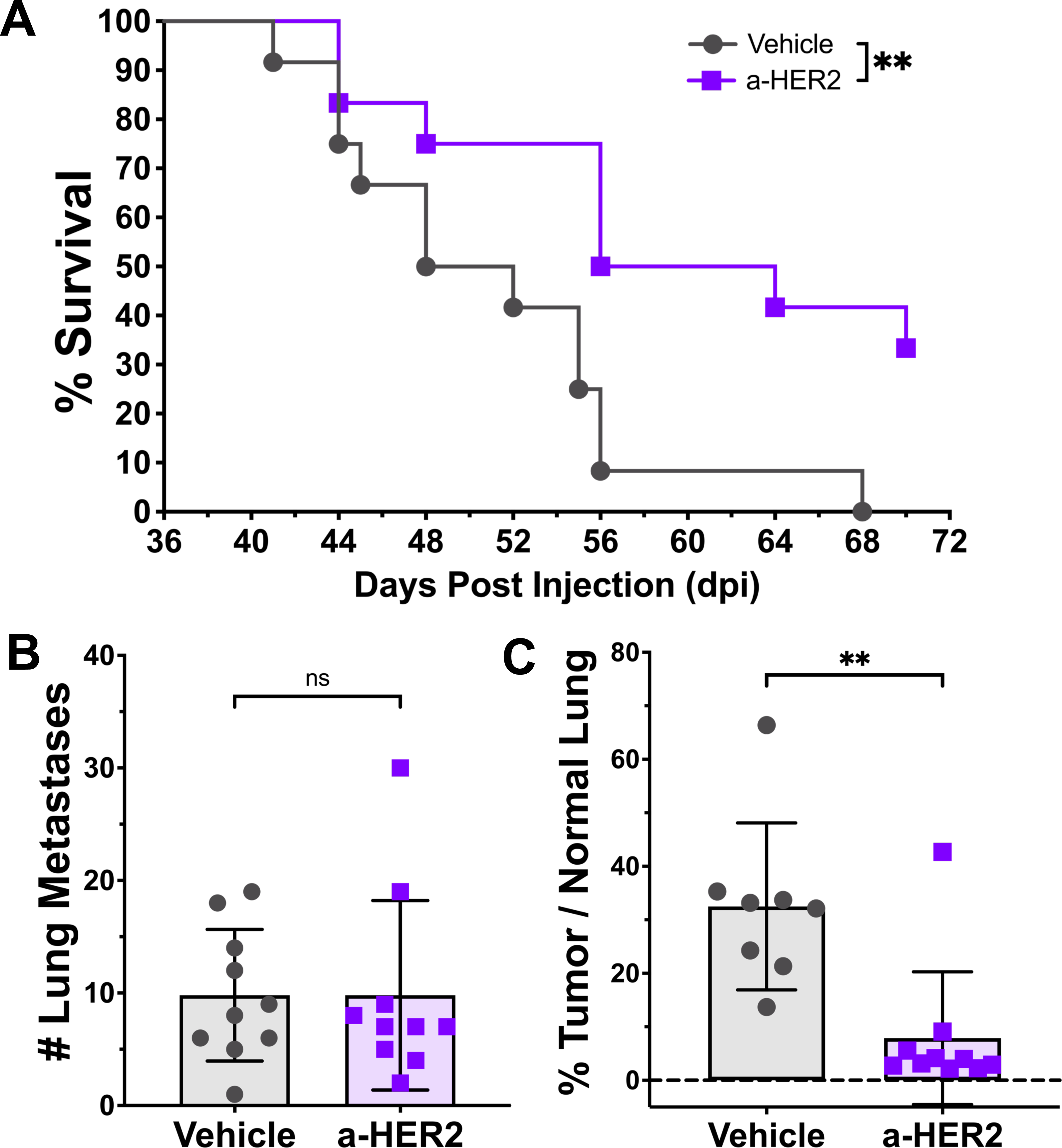
NT2.5-LM responds to HER2-directed therapy. **(A)** 1x10^5^ NT2.5-LM cells were injected into the mammary fat pad of NeuN mice. After surgical resection of NT2.5-LM tumor-bearing mice at 12 days post-injection (dpi), treatment with vehicle or anti-HER2 monoclonal antibody (100 µg/mouse, 1x/week, intraperitoneal injection) began at 23 dpi (n=12 per treatment group) and continued until survival endpoint at 70 dpi. (**B)** 1x10^5^ NT2.5-LM cells were injected into the mammary fat pad of NeuN mice, tumors were surgically resected at 12 dpi, and anti-HER2 treatment (100 µg/mouse, 1x/week, intraperitoneal injection) began at 23 dpi (n=10 per treatment group). Lungs were collected at 38 dpi. Three different levels were taken from formalin-fixed and paraffin-embedded lungs sectioned 100 µm apart. Slides were H&E stained, scanned, and analyzed using HALO to obtain summed lung metastasis counts and **(C)** percent tumor area over normal lung tissue. Two mice in the vehicle group were removed due to inconsistencies between HALO results and physical examination of H&E slides. Statistics used: Mantel-Cox Log-rank test for (A), Mann-Whitney U-test for (B-C), ns = not-significant, **p < 0.01.

### NT2.5-LM does not exhibit altered mutational landscape compared to parental NT2.5

With the increased number of lung metastases in NT2.5-LM model, we hypothesized that there might be differences in the genomic landscape and pathogenic mutational burden between the NT2.5 and NT2.5-LM tumors. First, we performed whole exome sequencing on the NT2.5 and NT2.5-LM cell lines to identify potential variations in genes with known pathogenic mutations and in genes known to affect proliferation and metastasis. Many pathogenic gene mutations common to breast cancer(Gil Del Alcazar et al., 2022), such as *Pten, Brca2, Atm, Cdh1, Chek2, Nf1, Arid1a, Pik3ca, and Esr1*, revealed no alterations between NT2.5 and NT2.5-LM (**Fig. 3A**). Of note, NT2.5-LM contained mutations in *Brca1* and NT2.5 contained mutations in *Rad51c*, but both were found within intron regions, thus not affecting protein sequence. Since NT2.5-LM is a HER2^+^ cell line, we examined the *Erbb2* transcript sequence across both cell lines more thoroughly and found six mutations within the protein coding sequence. However, all six mutations were silent (**Fig. 3B**). Lastly, we assessed tumor mutational burden, given that it represents another factor that could affect response to therapy. We found 11.45 mutations per megabase in the NT2.5 and 13.45 mutations per megabase in the NT2.5-LM models, with similar distributions of high missense mutations, single nucleotide polymorphisms (SNPs), and tyrosine-to-cytosine and cytosine-to-tyrosine mutations (**Figs. 3C-D**). Collectively, these data suggest that phenotypic differences between the NT2.5 and NT2.5-LM models are not the result of diversified mutational burden in NT2.5-LM.

**Figure 3:**
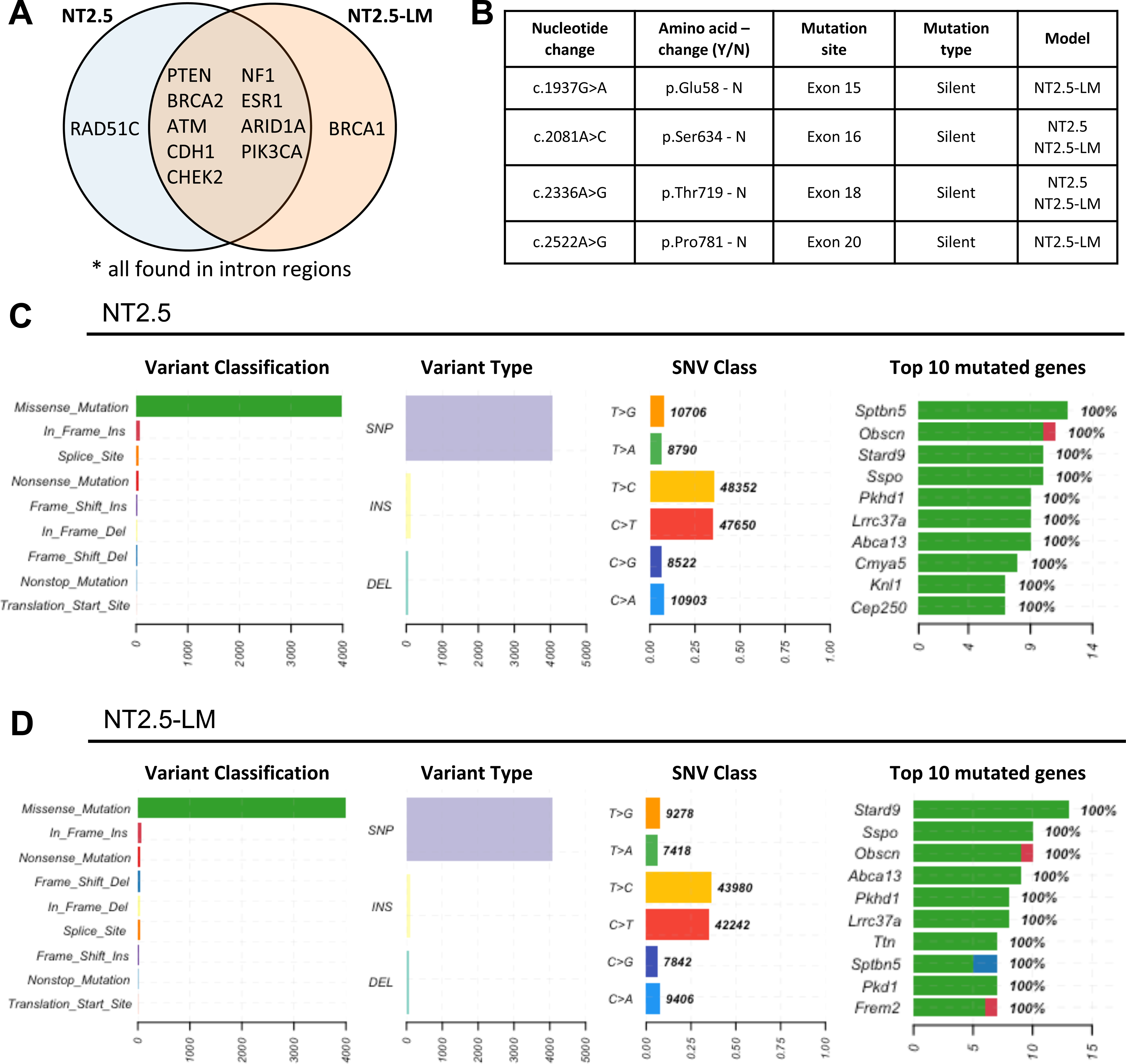
NT2.5-LM does not exhibit altered mutational landscape compared to parental NT2.5. **(A)** Alignment of NT2.5 and NT2.5-LM whole exome sequencing reads to the mm10 genome reveal cell line-specific and –overlapping mutations common in breast cancer. **(B)** Erbb2 transcript sequence with identified mutation sites in NT2.5 and NT2.5-LM. All mutations were identified to be silent mutations. Nucleotide numbering is based on DNA reference sequence NM_001003817.1. Note that the version number of this reference sequence may be frequently updated. **(C)** Distributions of mutation classifications, variant types, single nucleotide variant (SNV) classes, and top 10 mutated genes for NT2.5 and **(D)** NT2.5-LM are shown.

### NT2.5-LM exhibits altered signaling indicative of epithelial-to-mesenchymal transition (EMT)

Given the non-significant alterations in mutational burden, we sought to explain the differences in pro-metastatic phenotypes by comparing gene expression profiles between NT2.5 and NT2.5-LM. Four NT2.5 tumors and four NT2.5-LM tumors were collected from NeuN mice and subjected to unsorted single-cell RNA sequencing (scRNAseq), yielding approximately 9.6x10^8^ total reads. From Louvain clustering, approximately 10,000 NT2.5 and 9,000 NT2.5-LM cancer cells were identified as *Lcn+, Wfd2c+, Cd24a+, Cd276+, Col9a1+, Erbb2+,*(Berger et al., 2010; Gündüz et al., 2016; Seaman et al., 2017; Sidiropoulos et al., 2022; Yang et al., 2009; Yeo et al., 2020) subsetted out, and visualized by Principal Component Analysis (PCA) (**Fig. 4A**). An analysis of the top 25 differentially expressed genes between the two cancer cell clusters revealed an upregulation of genes associated with increased cellular proliferation [*Pdgfa*, *Sox9*],(Jansson et al., 2018; Ma et al., 2020; Pinto et al., 2014) invasion and migration [*Lrp1, Cd9, Cxcl1, Anxa1*],(Fayard et al., 2009; Moraes et al., 2018; Rappa et al., 2015; Xing et al., 2016; Yang et al., 2019) epithelial-to-mesenchymal transition (EMT) [*Vim, Inhba*], (Paulin et al., 2022; Yu et al., 2021) and stemness and metastatic potential [*S100A4, Nrp2, Aldh2, JunB*](Elaimy et al., 2018; Helfman et al., 2005; Qiao et al., 2015; Sundqvist et al., 2018; Yasuoka et al., 2009; Zhang and Fu, 2021) in NT2.5-LM. Concurrently, there was a downregulation of genes associated with decreased cellular proliferation [*Crip1*],(Ludyga et al., 2013) decreased invasion [*Cldn7*],(Kominsky et al., 2003; Martin and Jiang, 2009) and decreased epithelial phenotype and polarization [*Epcam*](Kyung-A Hyun et al., 2016) in NT2.5-LM (**Figs. 4B-C**). We validated the increased gene expression of *Vim* and decreased gene expression of *Epcam* in NT2.5-LM at the protein level by flow cytometry, demonstrating a significant increase in the percentage of Vimentin-positive cells and significant decrease in the percentage of Epcam-positive cells. (**Figs. 4D-E**).

**Figure 4:**
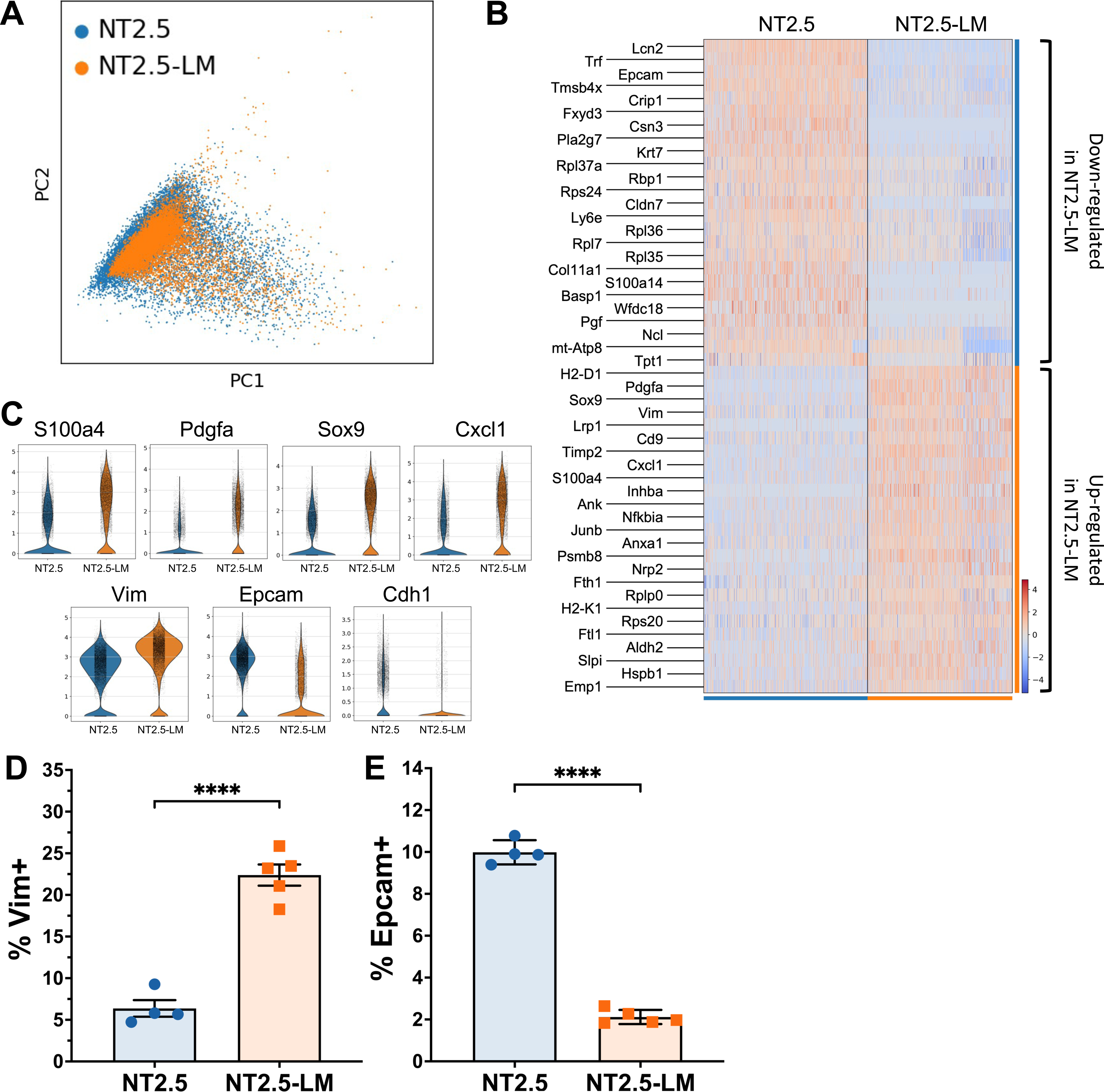
NT2.5-LM exhibits altered signaling indicative of increased EMT. **(A**) Four NT2.5 and four NT2.5-LM mammary tumors were collected from NeuN mice, dissociated to single cell suspensions, and sent for unsorted single-cell RNA sequencing. Cancer cell clusters were annotated as *Lcn+, Wfd2c+, Cd24a+, Cd276+, Col9a1+, Erbb2+,* and subsetted out for PCA visualization. (**B)** Top 25 significantly up- and down-regulated genes in NT2.5-LM. **(C)** Violin plots of key metastasis-related genes identified in (B). **(D)** Flow cytometry staining of epithelial-to-mesenchymal transition (EMT) related genes identified in (C) in NT2.5 and NT2.5-LM cell lines for Vimentin and **(E)** Epcam. Statistics used: Unpaired T-test for (D-E), ****p < 0.0001.

Further investigation into differential pathway regulation was performed by comparing the top 250 differentially expressed genes for overlap with pathways from the ‘KEGG_2019_Mouse’ database using Gene Set Enrichment Analysis. NT2.5-LM exhibited significant upregulation of the glycolysis pathway and downregulation of oxidative phosphorylation, ECM-receptor interaction, focal adhesion, protein digestion and absorption, and adherens junction pathways (p-adj < 0.05) (**Fig. S5, Table S1**). Dissolution of adherens junctions and alterations in cell-cell interactions is a hallmark of EMT,(Kalluri and Weinberg, 2009; Liu et al., 2016) and these data offer increased EMT as an explanation for the increased metastatic phenotype of NT2.5-LM.

### NT2.5-LM expresses increased levels of Mena^INV^ – a marker of metastatic potential

Our group has performed extensive work on mechanisms of metastatic dissemination and has previously reported that pro-migratory/pro-invasive tumor cells primed for the metastatic journey tend to upregulate the expression of Mena^INV^, a spliced isoform of the actin-regulatory protein mammalian enabled (Mena) that conveys increased metastatic potential. Specifically, previous studies have collectively shown that Mena^INV^ is correlated with increased breast cancer cell migration, invasion, and metastasis,(Borriello et al., 2022; Karagiannis et al., 2016; Philippar et al., 2008; Roussos et al., 2011b; Sharma et al., 2021) and is significantly upregulated in response to cytotoxic treatments.(Karagiannis et al., 2017) In view of observed alterations in various ECM and cell-cell adhesion interaction pathways, (**Fig. S5, Table S1**), we expected an enrichment of Mena^INV^-positive tumor cells in NT2.5-LM metastatic tumors. Indeed, immunofluorescence analysis of Mena^INV^ revealed significantly increased expression in the metastatic NT2.5-LM tumors, when compared to the NT2.5 mammary tumors (**Figs. 5A-B**).

**Figure 5:**
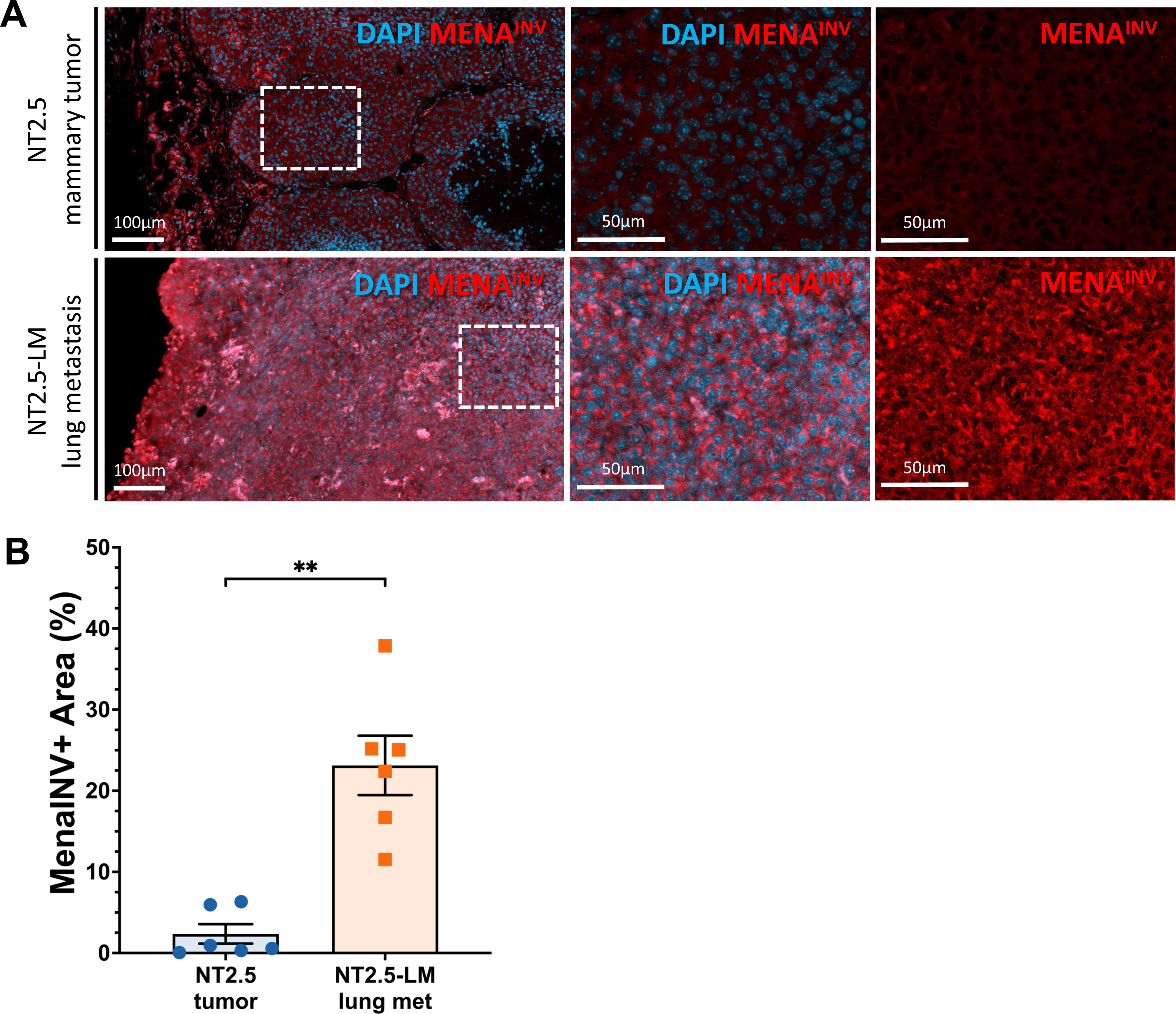
NT2.5-LM expresses increased levels of Mena^INV^ – a marker of metastatic potential. **(A**) Representative immunofluorescence images of Mena^INV^ (red) and DAPI (blue) staining in NT2.5 mammary tumor (top), and NT2.5-LM lung metastases (bottom) collected 34-41 days post-injection (dpi). Middle column and right column panels correspond to dotted square in left column panels. Scale bars as shown. **(B)** Quantification of Mena^INV^ staining from NT2.5 mammary tumor (n=6) and NT2.5-LM lung metastases (n=6) by averaging signal intensity from up to 10 regions of interest (ROIs) in each sample. Statistics used: Mann-Whitney U-test for (B), **p < 0.01.

## DISCUSSION

Spontaneously metastatic breast cancer cell lines are valuable tools for studying how metastatic tumors differ from primary tissue tumors in mice, but the time for spontaneous lung metastases to develop after injection of cancer cells into the breast tissue site is prolonged and inconsistent. In this study, we generated a more aggressively metastatic breast cancer cell line, NT2.5-LM, that spontaneously metastasizes to distant organs as early as one week post-injection. This not only allows us to study the effects of treatment interventions on metastatic progression in the most biologically accurate setting, but also utilizes surgical removal of the primary tumor early on to ensure that we are not limited by humane endpoints of primary tumor growth.

NT2.5-LM exhibited poorer survival, faster primary tumor growth, and more widespread metastases. Because the NT2.5-LM cell line was derived from NT2.5, we sought to understand the differences that would cause it to be more widely metastatic and proliferative compared to the parental cell line. We hypothesized that increased expression of HER2 or a novel mutation in the *ErbB2 gene* could be driving increased proliferation. NT2.5-LM did not exhibit new pathogenic mutations in *ErbB2,* and increased expression of HER2 was not observed by immunohistochemistry. Furthermore, pathways analyses conducted on scRNAseq data demonstrated no significant difference in expression of genes within the ErbB pathway. Thus, change in HER2 signaling is not a likely mechanism driving the increased metastatic and proliferative phenotype observed in NT2.5-LM.

Other potential mechanisms driving observed differences in NT2.5-LM include the differential regulation of proliferation- and metastasis-promoting pathways. We observed a shift in metabolic pathways with an upregulation of glycolysis and a downregulation of oxidative phosphorylation KEGG pathways, which have been previously implicated in more metastatic cancers,(Ashton et al., 2018; Gaude and Frezza, 2016) supporting our observations that NT2.5-LM is more widely metastatic. We observed a downregulation of ECM receptor interaction, focal junction, and adheres junction pathways, which are interactors in the intravasation and extravasation processes of metastasis.(Fares et al., 2020) We also identified differential expression of key genes involved in EMT that favored a more mesenchymal phenotype in NT2.5-LM, which could explain the increased number of metastases in lung and other distant organs. Our observed alterations in expression of epithelial markers, mesenchymal markers, cell adhesion pathways, extracellular matrix pathways, and metabolic pathways are characteristic of EMT.(Le Bras et al., 2012; Pal et al., 2022)

One interesting alteration associated with the loss of epithelial cell-cell contacts is the increased expression of invasive actin regulatory protein isoform Mena^INV^.(Goswami et al., 2009) Mena^INV^-expressing breast cancer cells participate in a paracrine loop with intratumoral macrophages, which facilitates their translocation to the perivascular niche. Once they reach the vasculature, Mena^INV^-expressing tumor cells associate with perivascular macrophages to intravasate into the blood vessel. These tripartite microanatomical structures composed of endothelial cells, perivascular macrophages, and Mena^INV^-expressing tumor cells are key prerequisites of metastatic dissemination and have been previously called Tumor Microenvironment of Metastasis (TMEM) doorways.(Borriello et al., 2022; Karagiannis et al., 2017; Philippar et al., 2008; Robinson et al., 2009; Roussos et al., 2011a; Sharma et al., 2021) Of note, NT2.5-LM tumors exhibit increased expression of Mena^INV^, which could explain its highly metastatic nature. As such, this model may be efficiently used in the future to study mechanisms of breast cancer cell dissemination associated with TMEM doorways and Mena^INV^-dependent pathways.

In summary, our findings distinguish NT2.5-LM as a more proliferative and metastatic model of breast cancer for experimental use that also preserves the spontaneous metastatic process within a shorter timeline. Various genetic and epigenetic changes can occur in a cancer cell as it accumulates mutations, proceeds through EMT, interacts with the TME, and forms distant metastases. Our group and others have shown that the addition of epigenetic modulators to various therapies in multiple cancer models has decreased tumor growth and improved response.(Brian J. Christmas et al., 2018; Kim et al., 2014; Orillion et al., 2017; Sidiropoulos et al., 2022) Moving forward, we envision the use of this NT2.5-LM model to facilitate efficient future studies of novel treatment combinations for metastatic disease and evaluation of different metastatic TME contributions to therapeutic response.

## METHODS

### Cell lines

NT2.5-lung metastasis (-LM) cell line was derived from the parental NT2.5 cell line, which was originally derived from the NT2 cell line in the NeuN murine model established by Guy et al.(C T Guy et al., 1992) 1x10^5^ NT2.5 cells were injected intravenously by tail vein in five 8-week-old female NeuN mice. Three weeks after tail vein injection, lung metastases were macro-dissected from all mice, minced on ice, filtered using a 100 µm filter, and pooled. The pooled cells were used to repeat the process described above, starting with intravenous injection, and after the third round of lung metastasis harvest, pooled cells were injected into the mammary fat pad of five 8-week-old female NeuN mice for spontaneous lung metastasis formation. After confirmation of spontaneous lung metastasis formation by lung harvest and Hematoxylin and Eosin (H&E) stains, the cell line was propagated in cell culture and named NT2.5-LM. NT2.5 cells were derived from spontaneous mammary tumors growing in female NeuN mice and obtained from the Jaffee Lab at Johns Hopkins University.(Jaffee et al., 1998; Machiels et al., 2001; R T Reilly et al., 2000b) Culture conditions for NT2.5-LM and NT2.5 cells are as follows: 37°C, 5% CO_2_ in RPMI 1640 (Gibco, cat. 11875-093) supplemented with 20% fetal bovine serum (Gemini, cat. 100-106), 1.2% HEPES (Gibco, cat. 15630-080), 1% L-glutamine (Gibco, cat. 25030-081), 1% MEM non-essential amino acids (Gibco, cat. 11140-050), 0.5% penicillin/streptomycin (Gibco, cat. 15140-122), 1% sodium pyruvate (Sigma, cat. S8636), 0.2% insulin (NovoLog, cat. U-100). Cell lines are tested for mycoplasma every 6 months.

### Mice

A syngeneic mouse model of HER2^+^ breast cancer using the NT2.5 cell line was derived from the NeuN transgenic mouse developed by Guy et al.(C T Guy et al., 1992) NeuN transgenic mice overexpress non-transforming rat neu cDNA under the control of a mammary specific promoter and develop spontaneous focal mammary adenocarcinomas after a long latency of 125 days with the majority of mice developing tumors by 300 days. Injection of NT2.5 into NeuN mice leads to development of tumors 100% of the time, since these mice are tolerized to Neu. Mice were kept in pathogen-free conditions and were treated in accordance with institutional and American Association of Laboratory Animal Committee policies. NeuN mice were originally from W. Muller McMaster University, Hamilton, Ontario, Canada and overexpress HER2 via the mouse mammary tumor virus (MMTV) promoter. Colonies are renewed yearly from Jackson labs and bred in-house by brother/sister mating.

### Survival, tumor growth, metastasis growth, necropsy

1x10^5^ NT2.5 or NT2.5-LM cells were injected into the mammary fat pad. NT2.5-LM tumors were resected on day 12. Survival endpoint was determined to be mammary tumor volume exceeding 1.5 cm^3^ or morbidity symptoms due to lung metastatic tumor burden, such as breathing, coat condition, activity, and posture. Mammary tumor growth was measured by calipers (± 0.01 mm) three times a week, with weekly tumor growth determined by calculating the average of differences in tumor volumes per week for each mouse. Lung surface metastases were counted by visual inspection of collected lungs following euthanasia at survival endpoint and before fixation in formalin and paraffin-embedding. Lung sections were taken 40 µm apart, for a representative 3 sections per lung. H&E stained sections were scanned and analyzed using either HALO or NDPView.2 to quantify number and tumor area of lung metastases. For necropsy, various tissues were collected at survival endpoint, fixed in formalin, paraffin-embedded, sectioned, stained with H&E, and visualized by light microscopy. Necropsy tissues include heart, lymph nodes, lungs, kidney, adrenal gland, stomach, colon, spleen, skull, ear, body wall, and teeth.

### Immunohistochemistry

Immunohistochemistry staining was performed at the Oncology Tissue Services Core of Johns Hopkins University. Immunolabeling for ErbB2, Ki67, CK5, CK6, AE1/3 and EGFR was performed on formalin-fixed, paraffin-embedded sections. Briefly, following dewaxing and rehydration, slides were immersed in 1% Tween-20, then heat-induced antigen retrieval was performed in a steamer using Antigen Unmasking Solution (catalog# H-3300, Vector Labs) for 25 minutes. Slides were rinsed in PBST, endogenous peroxidase and phosphatase were blocked (Dako, cat. S2003), and then incubated with the following primary antibodies for 45 minutes at room temperature: anti-ErbB2 (1:400 dilution; ThermoFisher Scientific, cat. MA5-15050, SF23975824), anti-Ki67 (1:200 dilution; Abcam, cat. Ab16667), anti-EGFR (1:50 dilution; LSBio, cat. LS-B2914-50), anti-CK5 (1:2000 dilution; BioLegend, cat. 905501), anti-CK6 (1:200 dilution; Novus Biologicals, cat. NBP2-34358), anti-AE-1/AE-3 (1:200 dilution; Novus Biologicals, cat. NBP2-29429). Slides were then incubated with HRP-conjugated anti-rabbit secondary antibody (Leica Microsystems, cat. PB6119) for 30 minutes at room temperature. Signal detection was conducted with 3,3′-Diaminobenzidine (Sigma-Aldrich, cat. D4293). Counterstaining was conducted with Mayer’s hematoxylin.

### Anti-HER2 treatment of mice

1x10^5^ NT2.5-LM cells were injected into the mammary fat pad. Mammary tumors were resected on day 12, after which mice were treated with anti-HER2 antibody starting on day 23 to mimic standard therapy treatment with trastuzumab in patients with HER2^+^ breast cancer. Anti-HER2 monoclonal antibody (BioXCell, clone 7.16.4) and mouse IgG2a isotype vehicle antibody (BioXCell, clone C1.18.4) were administered at 100 µg/mouse by intraperitoneal (i.p.) injection once a week for three weeks as described.(Brian J Christmas et al., 2018) Following three weeks of treatment, either lung tissues were collected for tumor burden analysis, or maintenance dosing was continued once a week until survival endpoint. For tumor burden analysis, three different levels were taken from formalin-fixed and paraffin-embedded lungs sectioned 100 µm apart. Slides were H&E stained, scanned, and analyzed using HALO to obtain summed lung metastasis counts and percent tumor area.

### Tumor dissociation

Following collection, mammary tumors were minced on ice and dissociated using a tumor dissociation kit (Miltenyi Biotec, cat. 130-096-730) and the 37C_m_TDK_2 program on the OctoDissociator (Miltenyi Biotec) per the manufacturer’s instructions. Cell suspensions were filtered using 70 µm cell strainers and red blood cells were lysed using ACK lysis buffer (Quality Biological, cat. 118-156-721). To submit for RNA sequencing, dead cells were removed using the MACS Dead Cell Removal Kit (Miltenyi Biotec).

### Flow cytometry

NT2.5 and NT2.5-LM cells were cultured for at least two passages, washed with PBS, and stained with Live/Dead Fixable Aqua (ThermoFisher, cat. L10119) for 30 minutes at 4°C, per the manufacturer’s instructions. Cells were fixed and permeabilized for 30 minutes at room temperature using the Foxp3 / Transcription Factor Staining Buffer Set (Life Technologies Corp., cat. 00-5523-00), followed by an Fc receptor block (BD Pharmingen, cat. 553142) for 10 minutes at room temperature. Cells were incubated with the following primary antibodies for 30 minutes at room temperature: anti-Vimentin (1:100 dilution; Cell Signaling Technology, cat. 5741), anti-Epcam (1:100 dilution; Cell Signaling Technology, cat. 93790). Cells were then incubated with FITC-conjugated anti-rabbit secondary antibody (1 µg/mL; BioLegend, cat. 406403) for 30 minutes at room temperature. Samples were run on the Attune NxT flow cytometer (Invitrogen) and analyzed using Kaluza software.

### Mena^INV^ Immunofluorescence and Image Analysis

Immunofluorescence staining for Mena^INV^ was performed on formalin fixed, paraffin-embedded (FFPE) sections. Briefly, slides were deparaffinized by melting for 5 minutes at 58°C in an oven equipped with a fan, followed by two Xylene treatments for 20 minutes each. Slides were rehydrated and antigen retrieval was performed in 1 mM EDTA, pH 8.0 for 20 minutes at 97°C in a conventional steamer. Slides were washed with 0.05% PBST and incubated in blocking solution (5% goat serum in 0.05% PBST) for 1 hour at room temperature. Slides were then incubated with anti-Mena^INV^ primary antibody (0.25 ug/mL; in-house developed in the lab of Dr. John S. Condeelis, AE1071, AP-4) overnight at 4°C. After three washes in 0.05% PBST, slides were incubated with Alexa 488-conjugated goat anti-chicken secondary antibody at room temperature for 1 hour. After three washes in 0.05% PBST, slides were incubated with spectral DAPI for 5 minutes and mounted with ProLong Gold Antifade Mountant (Life Technologies, cat. P36930). Slides were imaged using the Pannoramic 250 Flash II digital whole slide scanner. Up to 10 High-Power Field (HPF) images per mouse, depending on tumor and metastasis burden availability, were captured in TIFF format using Caseviewer v2.4 (3DHISTECH). Further image processing was performed in ImageJ. Single Mena^INV^ channels were uploaded, converted to 8-bit, and binarized using intensity thresholding (default method). The DAPI channel confirmed that all HPFs chosen were within necrosis-free areas of the tumors and metastases. The Mena^INV+^ area in each HPF was then expressed as a fraction of the total tumor area, and the mean of all HPFs was calculated for each mouse. For visualization purposes only, images were enhanced in Caseviewer by exclusively using linear image modifications (i.e., brightness and contrast), and the signal was pseudo-colored for optimal representation of fields of interest.

### Whole exome sequencing (WES)

NT2.5 and NT2.5-LM cell lines were cultured as described above and sent for whole exome sequencing at the Johns Hopkins Genomics Core. One microgram or more of mouse genomic DNA from each sample was analyzed by whole exome sequencing using the SureSelectXT Mouse All Exon kit (Agilent), followed by next generation sequencing using the NovaSeq 6000 S4 flow cell (Illumina) with a 2x150bp paired-end read configuration, per the manufacturer’s instructions. bcl2fastq v2.15.0 (Illumina) was used to convert BCL files to FASTQ files using default parameters. Running alignments against the mm10 genome was done by bwa v0.7.7 (mem) along with Piccard-tools1.119 to add read groups and remove duplicate reads. GATK v3.6.0 base call recalibration steps were used to create a final alignment file. MuTect2 v3.6.0 was used to call somatic variants against a panel of normal using default parameters. snpEFF (v4.1) was used to annotate the variant calls and to create a clean tab separated table of variants. IGV v2.13.2 was used to identify breast cancer specific mutations from MuTect2 files. SnapGene Viewer v.6.2 was used to visually align and determine the mutations between the two cell lines against the mRNA sequences of selected genes. Annotations were created to visualize mutational differences.

### Single cell RNA sequencing (scRNA-seq)

For library preparation, 10x Genomics Chromium Single Cell 3′ RNA-seq kits v3 were used. Gene expression libraries were prepared per the manufacturer’s instructions. 4 biological replicates totaling 8 processed tumors were sequenced in 2 batches: Run A - 2 NT2.5 tumors, 2 NT2.5-LM tumors; Run B - 2 NT2.5 tumors, 2 NT2.5-LM tumors. These tumors were taken as a subset from a larger batch of tumors that include various mouse treatments, with each batch having an equal assortment of samples from multiple treatment groups to reduce technical biases. Here, we restrict our analysis to replicates under the vehicle treatment condition. Illumina HiSeqX Ten or NovaSeq were used to generate total reads. Paired-end reads were processed using CellRanger v3.0.2 and mapped to the mm10 transcriptome with default settings. ScanPy v1.8.2 and Python v3 was used for quality control and basic filtering. DoubleDetection v4.2 with Louvain clustering algorithm v0.7.1 was used to find doublets. For gene filtering, all genes expressed in less than 3 cells within a tumor (NT2.5 and NT2.5-LM) were removed. Cells expressing less than 200 genes or more than 8,000 genes or having more than 15% mitochondrial gene expression were also removed. Gene expression was total-count normalized to 10,000 reads per cell and log transformed. Highly variable genes were identified using default ScanPy parameters, and the total counts per cell and the percent mitochondrial genes expressed were regressed out. Finally, gene expression was scaled to unit variance and values exceeding 10 standard deviations were removed. Neighborhood graphs were constructed using 10 nearest neighbors and 30 principal components. Tumors were clustered together within cell lines using Louvain clustering (with resolution parameter 0.12) and cancer cells were identified as *Lcn+, Wfd2c+, Cd24a+, Cd276+, Col9a1+, Erbb2+.*(Berger et al., 2010; Gündüz et al., 2016; Seaman et al., 2017; Sidiropoulos et al., 2022; Yang et al., 2009; Yeo et al., 2020) All other cell clusters and doublets were removed. There were ∼10,000 NT2.5 cancer cells and ∼9,000 NT2.5-LM cancer cells, and these were combined by total raw count normalization to 10,000 reads, with log transformation and batch correction on cell lines via ComBat. The 250 top differentially expressed genes in the cancer clusters from each cell line were identified using the Wilcoxon rank-sum test and compared for overlap with pathways from the ‘KEGG_2019_Mouse’ database using GSEAPY (Gene Set Enrichment Analysis in Python).

### Statistics

For survival curves, Mantel-Cox log rank tests were used. For tumor growth rate, metastasis counts, and lung metastasis volumes, Mann Whitney tests were used. For quantification of immunohistochemistry staining, Welch’s T-tests were used. For flow cytometry, unpaired t-tests were used. For immunofluorescence staining of tumor and metastatic tissues, Mann Whitney U-tests were used. To aid in statistical choice, data were tested for normality using D’Agostino-Pearson omnibus normality tests, Anderson-Darling tests, Shapiro-Wilk normality tests, and Kolmogorov-Smirnov normality tests.

## DECLARATIONS

## Acknowledgements

We would like to thank all members of the Elizabeth Jaffee and Elana Fertig lab for help throughout the course of these experiments. Additionally, we would like to thank the Molecular Genomics Core at USC, the Flow Cytometry Core at USC, the Translational Pathology Adult Tissue Core at USC, the SKCCC Experimental and Computational Genomics Core at Johns Hopkins, and the Oncology Tissue Services Core at Johns Hopkins for help with sequencing experiments, specimen processing, and data processing. We would like to thank the Analytical Imaging Facility at the Albert Einstein College of Medicine for immunofluorescence and tissue slide scanning. We would also like to acknowledge Srinivasan Yegnasubramanian and Emma Bigelow from the Johns Hopkins research group for data review.

## Availability of data and materials

All WES and scRNAseq raw and processed data files will be made available on NCBI BioProject.

## Ethics approval and consent to participate

All animal studies were approved by the Institutional Review Board of USC and Johns Hopkins University.

---

**Figure S1: Tumor growth in NT2.5-LM model. (A)** 1x10^5^ NT2.5 or NT2.5-LM cells were injected into the mammary fat pad of NeuN mice (NT2.5, n=10; NT2.5-LM, n=7). Mammary tumor volumes (mm^3^) were averaged across all mice within the same group. Surgical resection of NT2.5-LM tumor-bearing mice at 12 days post-injection (dpi) is depicted by a red arrow. Mammary tumors regrew in NT2.5-LM at 24 dpi. Data shown until first mouse death recorded at 33 dpi. **(B)** Mammary tumor volumes (mm^3^) of individual mice shown in (A) until required euthanasia of mice.

**Figure S2: Necropsy of NT2.5-LM metastases-bearing tissues**. Upon euthanasia of NT2.5-LM mice, various tissues were collected, fixed, sectioned, stained with H&E, and evaluated for the presence of metastases. Tissues shown include (A) heart [scale bars: 1000 µm], (B) lymph nodes [scale bars: 50 µm, 1000 µm], (C) lungs [scale bar: 2500 µm], (D) kidney [scale bar: 500 µm], (E) adrenal gland [scale bar: 500 µm], (F) stomach [scale bars: 500 µm, 1000 µm], (G) colon [scale bars: 400 µm, 2500 µm], (H) spleen [scale bar: 250 µm], (I) skull [scale bar: 2500 µm], (J) ear [scale bar: 5000 µm], (K) body wall [scale bar: 2500 µm], and (L) teeth [scale bars: 50 µm, 750 µm].

**Figure S3: Immunohistochemistry (IHC) of NT2.5 mammary tumors and NT2.5-LM lung metastases.** Staining of EGFR, AE1/3, CK5, and CK6 in NT2.5 mammary tumors (left) and NT2.5-LM lung metastases (right) collected at 35 days post-injection. Scale bars are 280 µm and 60 µm (zoomed-in panels).

**Figure S4: Anti-HER2 treatment scheme for NT2.5-LM**. 1x10^5^ NT2.5-LM cells were orthotopically injected in the mammary fat pad. Mammary tumors were surgically resected 12 days post-injection (dpi). Anti-HER2 monoclonal antibody treatment of 100 µg/mouse administered intraperitoneally once a week for three weeks began at 23 dpi. After three weeks of anti-HER2 treatment, maintenance dosage for survival experiments were given once a week. For metastatic burden analysis, lungs were collected at 38 dpi for subsequent analysis.

**Figure S5: Differential pathway regulation in NT2.5-LM compared to NT2.5 cancer cells**. Unsupervised pathways analysis from single cell RNA sequencing datasets by comparing top 250 differentially expressed genes with overlap in pathways from ‘KEGG_2019_Mouse’ database using Gene Set Enrichment Analysis. Top 20 pathways in NT2.5-LM that are **(A)** down-regulated and **(B)** up-regulated compared to NT2.5 are shown.

**Table S1**: **Differential pathways in NT2.5-LM compared to NT2.5 cancer cells**. All unsupervised pathways analysis from single cell RNA sequencing datasets by comparing top 250 differentially expressed genes with overlap in pathways from ‘KEGG_2019_Mouse’ database using Gene Set Enrichment Analysis.

