## Supplemental Figures for "Mimicking the breast metastatic microenvironment: characterization of a novel syngeneic model of HER2^+^ breast cancer"

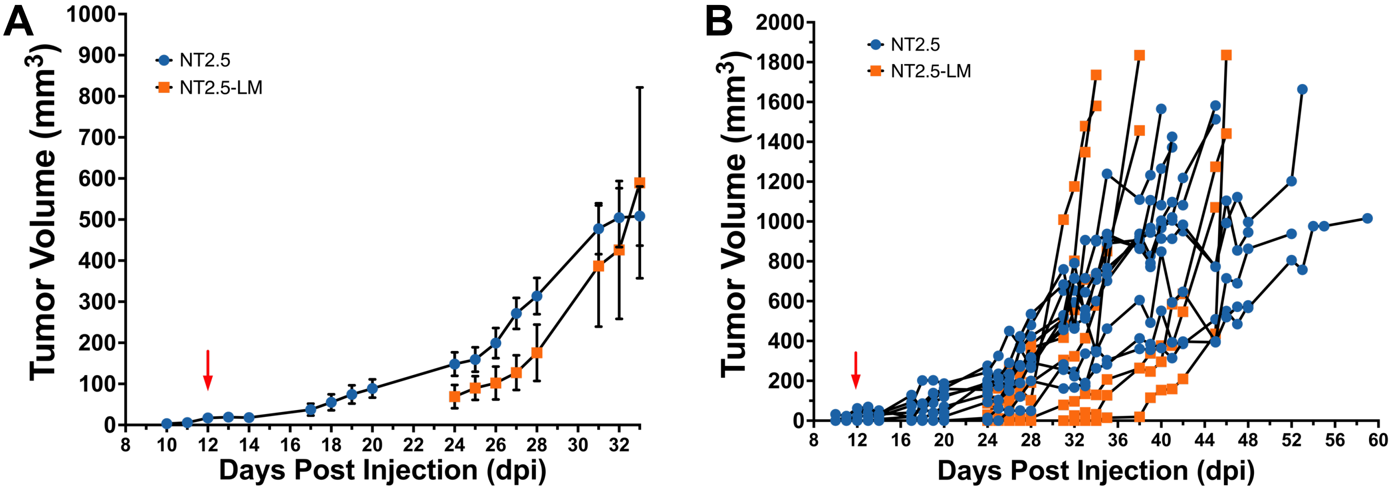

A
Tumor Volume (mm3)
NT2.5
NT2.5-LM
Days Post Injection (dpi)
B
Tumor Volume (mm3)
NT2.5
NT2.5-LM
Days Post Injection (dpi)


Figure S1

**Figure S1: Tumor growth in NT2.5-LM model. (A)** $1x10^5$ NT2.5 or NT2.5-LM cells were injected into the mammary fat pad of NeuN mice (NT2.5, n=10; NT2.5-LM, n=7). Mammary tumor volumes ($mm^3$) were averaged across all mice within the same group. Surgical resection of NT2.5-LM tumor-bearing mice at 12 days post-injection (dpi) is depicted by a red arrow. Mammary tumors regrew in NT2.5-LM at 24 dpi. Data shown until first mouse death recorded at 33 dpi. **(B)** Mammary tumor volumes ($mm^3$) of individual mice shown in (A) until required euthanasia of mice.

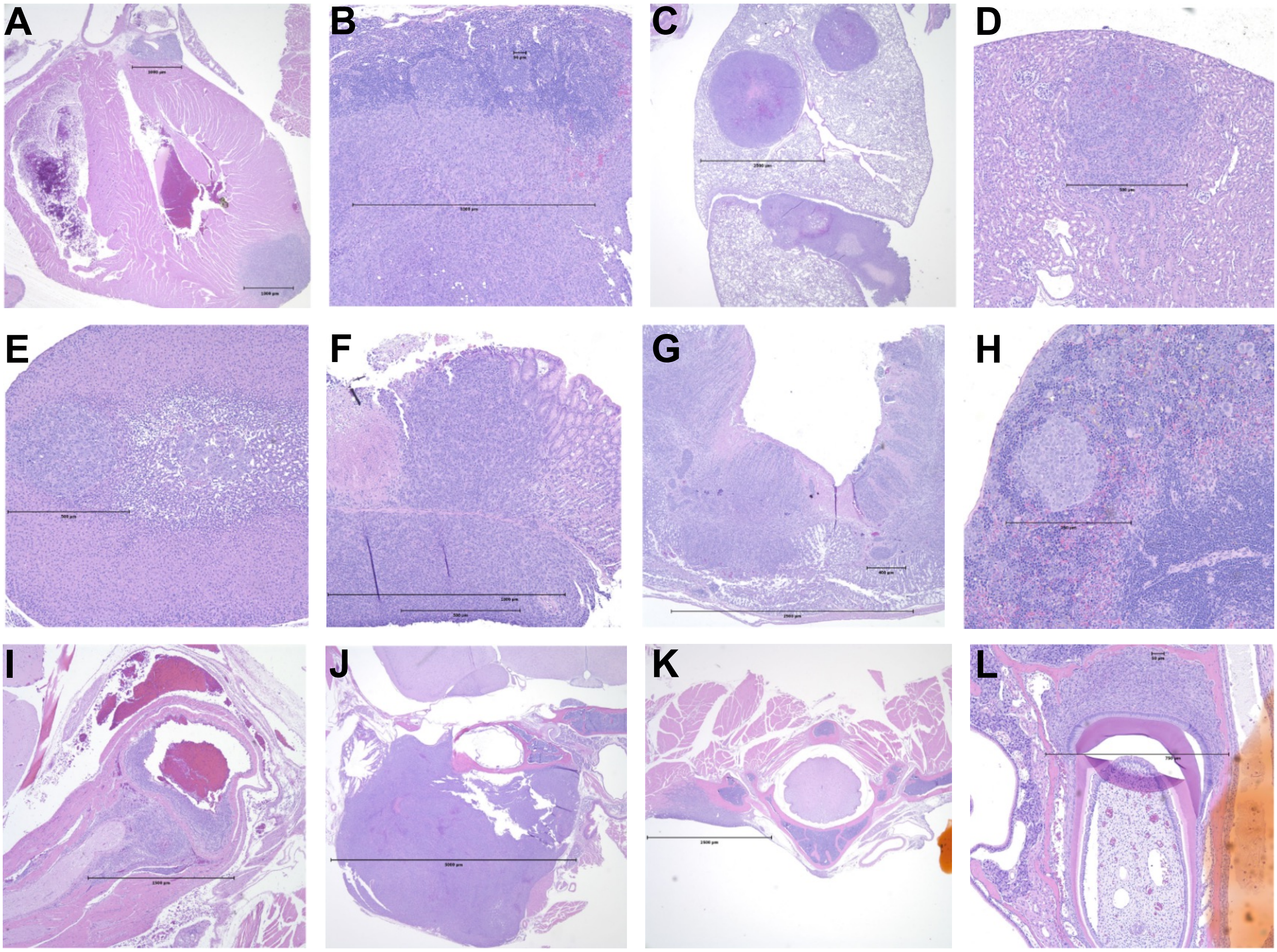


**Figure S2**

**Figure S2: Necropsy of NT2.5-LM metastases-bearing tissues**. Upon euthanasia of NT2.5-LM mice, various tissues were collected, fixed, sectioned, stained with H&E, and evaluated for the presence of metastases. Tissues shown include (A) heart [scale bars: 1000 μm], (B) lymph nodes [scale bars: 50 μm, 1000 μm], (C) lungs [scale bar: 2500 μm], (D) kidney [scale bar: 500 μm], (E) adrenal gland [scale bar: 500 μm], (F) stomach [scale bars: 500 μm, 1000 μm], (G) colon [scale bars: 400 μm, 2500 μm], (H) spleen [scale bar: 250 μm], (I) skull [scale bar: 2500 μm], (J) ear [scale bar: 5000 μm], (K) body wall [scale bar: 2500 μm], and (L) teeth [scale bars: 50 μm, 750 μm].

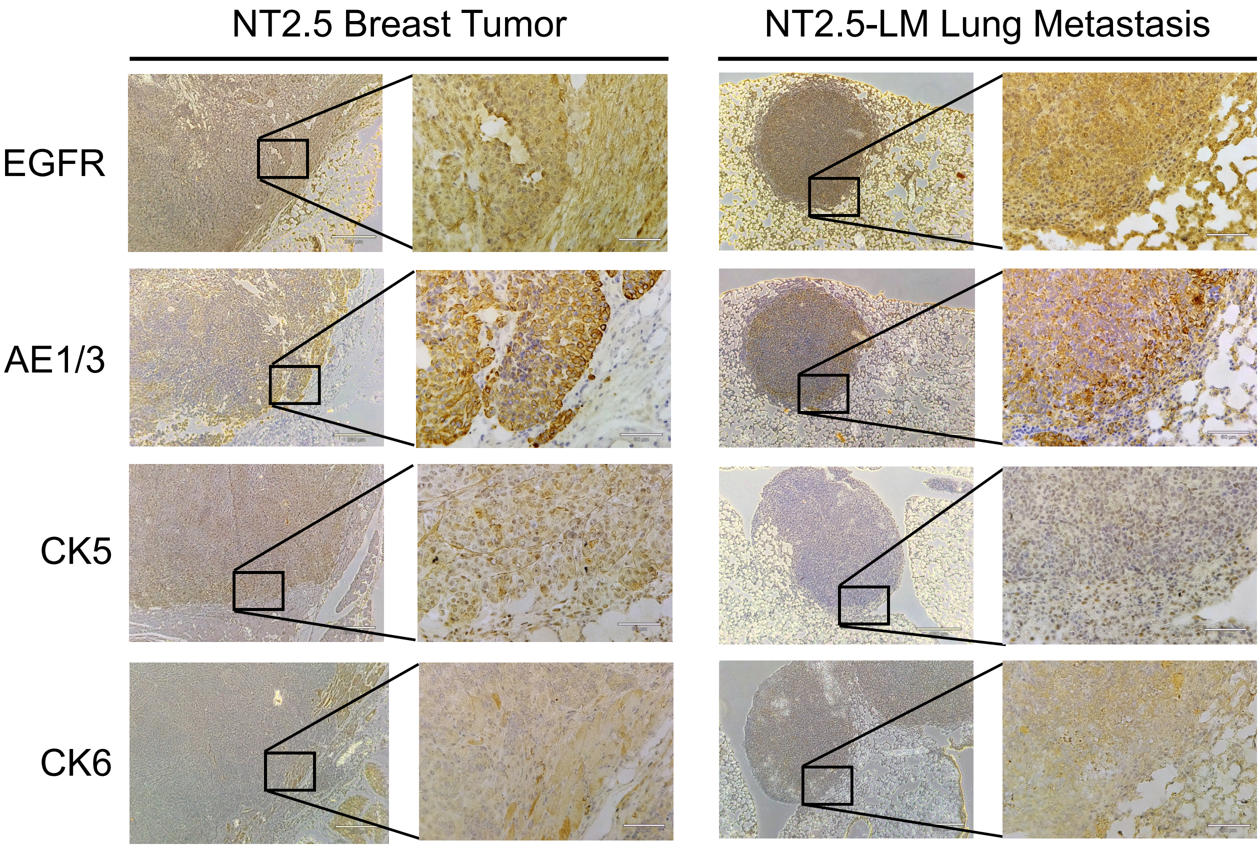


**Figure S3**

**Figure S3: Immunohistochemistry (IHC) of NT2.5 mammary tumors and NT2.5-LM lung metastases.** Staining of EGFR, AE1/3, CK5, and CK6 in NT2.5 mammary tumors (left) and NT2.5-LM lung metastases (right) collected at 35 days post-injection. Scale bars are 280 μm and 60 μm (zoomed-in panels).

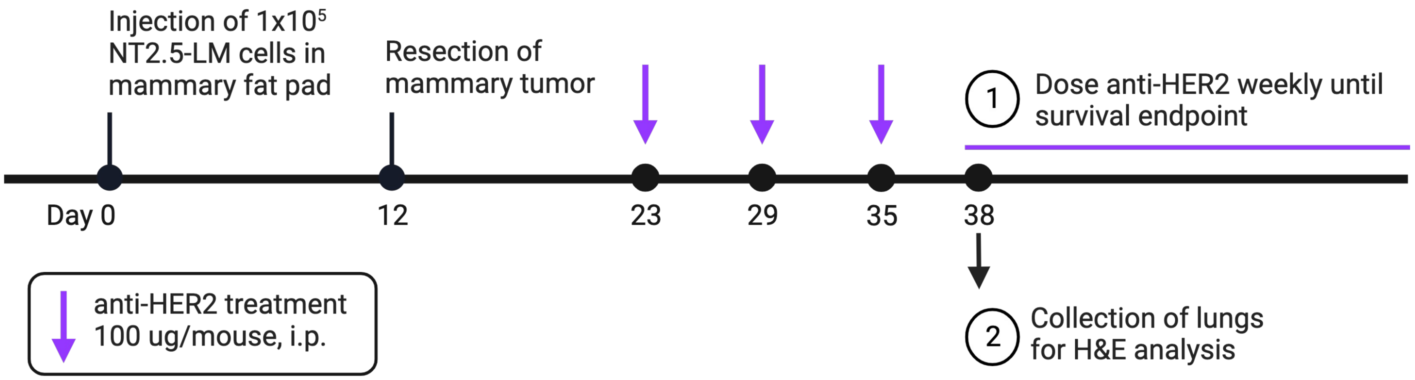
Injection of 1x10^5 NT2.5-LM cells in mammary fat pad
Resection of mammary tumor
1 Dose anti-HER2 weekly until survival endpoint
Day 0
12
23
29
35
38
anti-HER2 treatment 100 ug/mouse, i.p.
2 Collection of lungs for H&E analysis

**Figure S4**

**Figure S4: Anti-HER2 treatment scheme for NT2.5-LM**. $1x10^5$ NT2.5-LM cells were orthotopically injected in the mammary fat pad. Mammary tumors were surgically resected 12 days post-injection (dpi). Anti-HER2 monoclonal antibody treatment of 100 µg/mouse administered intraperitoneally once a week for three weeks began at 23 dpi. After three weeks of anti-HER2 treatment, maintenance dosage for survival experiments were given once a week. For metastatic burden analysis, lungs were collected at 38 dpi for subsequent analysis.

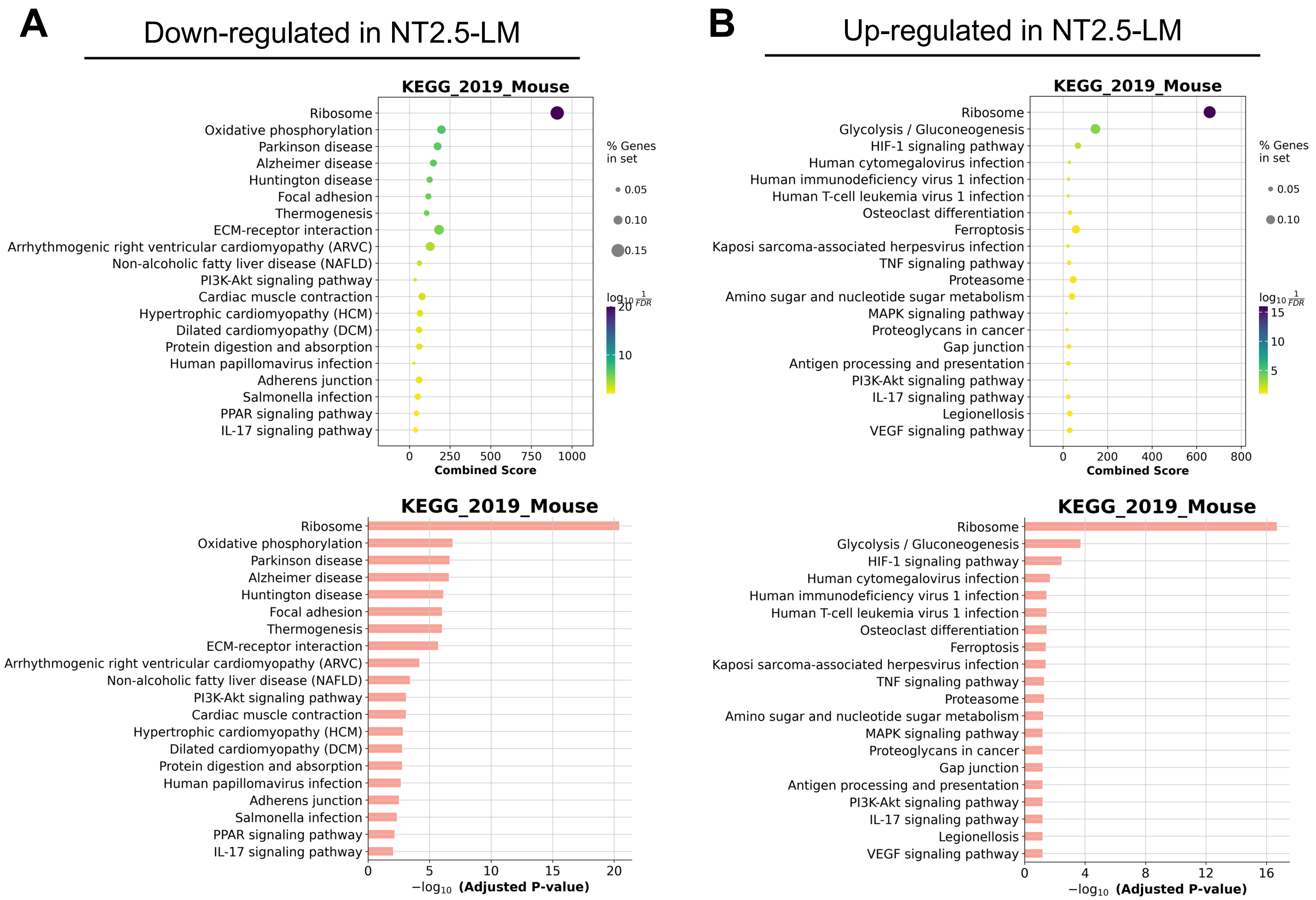
A
Down-regulated in NT2.5-LM
KEGG_2019_Mouse
Ribosome
Oxidative phosphorylation
Parkinson disease
Alzheimer disease
Huntington disease
Focal adhesion
Thermogenesis
ECM-receptor interaction
Arrhythmogenic right ventricular cardiomyopathy (ARVC)
Non-alcoholic fatty liver disease (NAFLD)
PI3K-Akt signaling pathway
Cardiac muscle contraction
Hypertrophic cardiomyopathy (HCM)
Dilated cardiomyopathy (DCM)
Protein digestion and absorption
Human papillomavirus infection
Adherens junction
Salmonella infection
PPAR signaling pathway
IL-17 signaling pathway
Combined Score
% Genes in set
0.05
0.10
0.15
$\log_{10}\frac{1}{FDR}$
−log10 (Adjusted P-value)
B
Up-regulated in NT2.5-LM
KEGG_2019_Mouse
Ribosome
Glycolysis / Gluconeogenesis
HIF-1 signaling pathway
Human cytomegalovirus infection
Human immunodeficiency virus 1 infection
Human T-cell leukemia virus 1 infection
Osteoclast differentiation
Ferroptosis
Kaposi sarcoma-associated herpesvirus infection
TNF signaling pathway
Proteasome
Amino sugar and nucleotide sugar metabolism
MAPK signaling pathway
Proteoglycans in cancer
Gap junction
Antigen processing and presentation
PI3K-Akt signaling pathway
IL-17 signaling pathway
Legionellosis
VEGF signaling pathway
Combined Score
% Genes in set
0.05
0.10
$\log_{10}\frac{1}{FDR}$
−log10 (Adjusted P-value)

**Figure S5**

**Figure S5: Differential pathway regulation in NT2.5-LM compared to NT2.5 cancer cells**. Unsupervised pathways analysis from single cell RNA sequencing datasets by comparing top 250 differentially expressed genes with overlap in pathways from 'KEGG_2019_Mouse' database using Gene Set Enrichment Analysis. Top 20 pathways in NT2.5-LM that are **(A)** down-regulated and **(B)** up-regulated compared to NT2.5 are shown.

**Table S1**: **Differential pathways in NT2.5-LM compared to NT2.5 cancer cells**. All unsupervised pathways analysis from single cell RNA sequencing datasets by comparing top 250 differentially expressed genes with overlap in pathways from 'KEGG_2019_Mouse' database using Gene Set Enrichment Analysis.
